# *Arabidopsis* HAP2/GCS1 is a gamete fusion protein homologous to somatic and viral fusogens

**DOI:** 10.1101/097568

**Authors:** Clari Valansi, David Moi, Evgenia Leikina, Elena Matveev, Martín Graña, Leonid V Chernomordik, Héctor Romero, Pablo S. Aguilar, Benjamin Podbilewicz

**Author notes:** These authors contributed equally to this work (Clari Valansi, David Moi).

## Abstract

Cell-cell fusion is inherent to any form of sexual reproduction. Loss of HAPLESS 2/GENERATIVE CELL SPECIFIC 1 (HAP2/GCS1) proteins results in gamete fusion failure in different organisms but their exact role is unclear. Here we show that *Arabidopsis* HAP2/GCS1 expression in mammalian cells is sufficient to promote cell-cell fusion. Hemifusion and complete fusion depend on HAP2/GCS1 presence in both fusing cells. Furthermore, expression of HAP2 on the surface of pseudotyped vesicular stomatitis virus and on the target cells results in HAP2-dependent virus-cell fusion. This bilateral requirement can be bypassed by replacing the plant gene with *C. elegans* EFF-1 somatic cell fusogen in one of the fusing cells or the virus, indicating that HAP2/GCS1 and EFF-1 share a similar fusion mechanism. Structural modeling of the HAP2/GCS1 protein family predicts that they are homologous to EFF-1 and class II fusion proteins from enveloped viruses (e.g. dengue and Zika viruses). We name this superfamily FUSEXINS: FUSion proteins essential for sexual reproduction and EXoplasmic merger of plasma membranes. Thus, Fusexins unify the origin and evolution of sexual reproduction, enveloped virus entry into cells and somatic cell fusion.

## Introduction

While proteins mediating cell-cell fusion in tissues have been demonstrated in the placenta of mammals (Syncytins) and in organs of invertebrates (e.g. EFF-1 in *C. elegans*), the machinery mediating sperm-egg fusion remains unknown (Aguilar et al., 2013; Bianchi et al., 2014; Maruyama et al., 2016). HAP2/GCS1 proteins have been implicated as potential gamete fusogens in *Arabidopsis* (Johnson et al., 2004; Mori et al., 2006; von Besser et al., 2006), *Chlamydomonas* (Liu et al., 2008), *Tetrahymena* (Cole et al., 2014), *Dictyostelium* (Okamoto et al., 2016) and *Plasmodium* (Liu et al., 2008). However the precise function of HAP2/GCS1 in gamete fusion is unknown. So far, there is no functional or structural evidence indicating HAP2/GCS1 is directly involved in cell-cell fusion. Proteins may function as direct fusogens, or alternatively, they may affect communication or intimate adhesion before fusion takes place, as demonstrated for other gamete fusion candidates such as Juno and Izumo receptors (Bianchi et al., 2014).

## Results and Discussion

To determine whether HAP2/GCS1 is an authentic fusion protein we first tested whether *Arabidopsis* HAP2 (AtHAP2) could fuse heterologous cells that normally do not fuse. For this, we transfected Baby Hamster Kidney (BHK) cells with plasmids encoding AtHAP2, EFF-1, or red (RFP) or green (GFP) fluorescent proteins as negative controls, and assayed the extent of cell-cell fusion (Fig. 1 A). In controls, when BHK cells were transfected with cytoplasmic RFP (RFPcyto-BHK) and mixed with GFP-transfected BHK cells (GFP-BHK; Fig. 1 B (i)) about 5% of cells (red or green, respectively) had two nuclei due to cell division and only 1.5% of the cells expressed both GFP and RFPcyto out of the total GFP/RFPcyto expressing cells in contact (Fig. 1 C). This apparent cytoplasmic content mixing could be due to phagocytosis of fluorescent apoptotic bodies or background fusion. In contrast, when AtHAP2 was transfected into BHK cells with either RFPcyto or GFP and the transfected cells were co-incubated, we observed an average multinucleation of 33% ± 3 and 41.3% ± 1.3 (green or red), and cytoplasmic content mixing in 11.3% ± 0.9 in three independent experiments (Fig. 1 B (ii, iv) and C). Similar results were obtained using the previously defined *C. elegans’* somatic cell fusogen EFF-1 (Fig. 1B (iii) and C). To test whether some of the multinucleated cells resulted from faulty cell division (e.g. nuclear division without cytokinesis) we incubated BHK-HAP2 cells with fluorodeoxyuridine (FdUrd) to arrest the cell cycle at the G1/S transition. We found that FdUrd failed to decrease multinucleation in HAP2-BHK cells, and actually doubled the multinucleation, probably due to a block in cell division of multinucleated cells formed by cell-cell fusion. In contrast, FdUrd did not significantly affect nuclei content of BHK cells that were transfected with the GFP plasmid only (Table S1). Time-lapse microscopy experiments showed that mononucleated BHK-HAP2(RFPcyto) cells formed syncytia by merging their cytoplasms (Fig. 1 D; **Videos 1-4; Fig. S1 A**). Thus, AtHAP2 expression in BHK cells is sufficient to promote cell-cell fusion, defining this protein as a bona fide fusogen.

**Figure 1.**
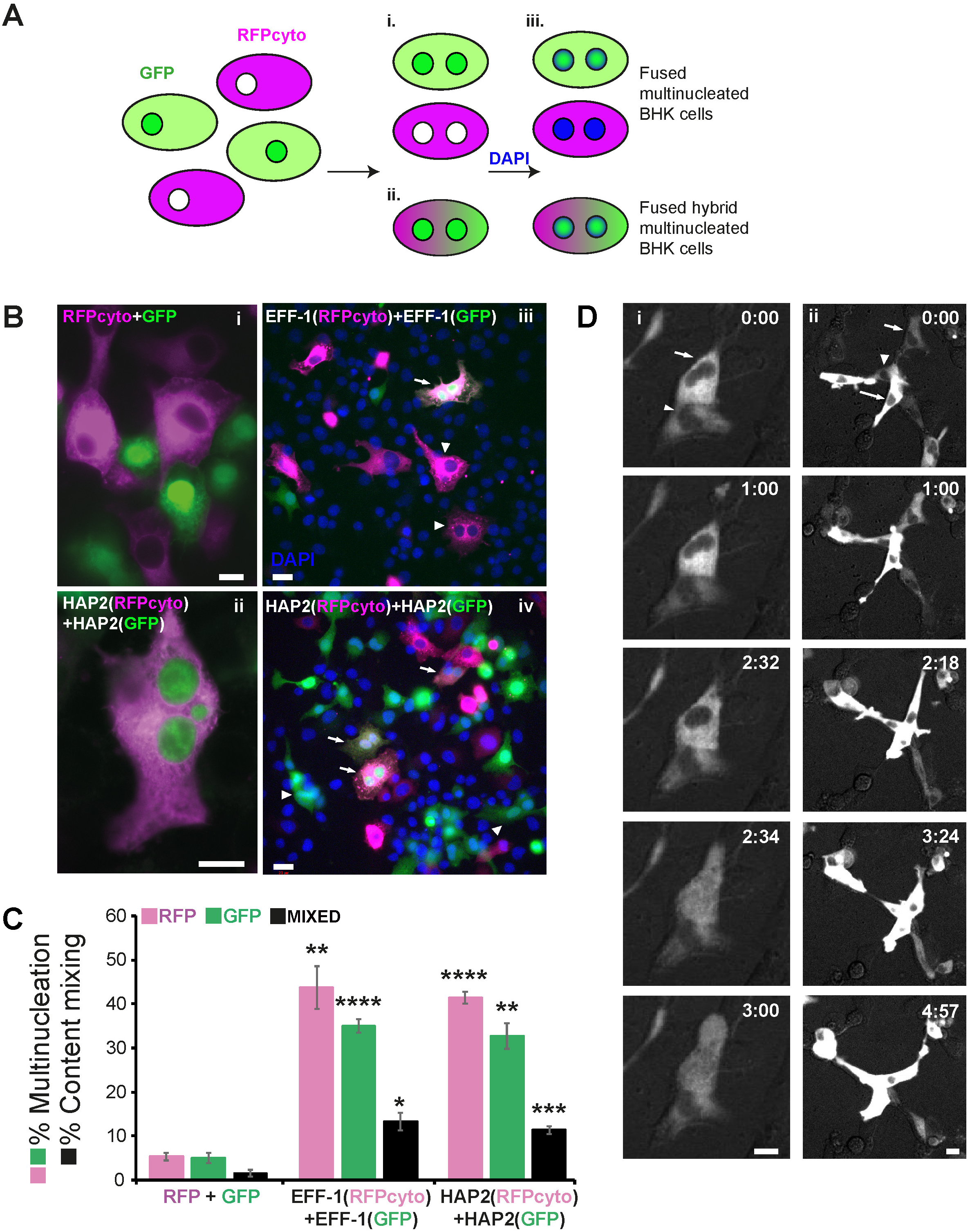
Arabidopsis HAP2 is sufficient to fuse mammalian BHK cells. **(A)** BHK cell-cell fusion assay: After discarding a possible failure in cell division (**Table S1**) cell-cell fusion is measured by the appearance of **i)** multinucleated cells labelled with either cytoplasmic RFP (RFPcyto; magenta) or nuclear and cytoplasmic GFP (green). Fusion is also indicated by the appearance **ii)** of multinucleated cells containing nuclear GFP and fluorescence from both RFPcyto and GFP in the cytoplasm. **iii)** Nuclei are labelled with DAPI (blue) after fixation and permeabilization of the cells. **(B) i) RFPcyto + GFP**: Negative control shows mononucleated cells expressing RFPcyto (magenta) or nuclear and cytoplasmic GFP (green). **ii) HAP2(RFPcyto) + HAP2(GFP):** BHK cells were transfected with AtHAP2 and GFP (green) or RFPcyto (magenta); merged image of hybrid cell that contains mixed cytoplasm and three nuclei. **iii) EFF-1(RFPcyto) + EFF-1(GFP):** Hybrid binucleate cell emerged after EFF-1 expression and mixing of magenta and green cells (arrow). EFF-1(RFPcyto) binucleate cells (arrowhead). **iv) HAP2(RFPcyto) + HAP2(GFP):** Heterokaryons (hybrids) express magenta cytoplasm and green nuclei and cytoplasm (arrows). Multinucleate green cells (arrowheads). Scale bars, i and ii 10 μm; (iii and iv) 20 μm. **(C)** Quantification of multinucleation and content mixing experiments. Magenta and green bars represent the fraction of multinucleated cells (two nuclei or higher) out of all the cells in contact (magenta or green respectively). Black bars represent the RFPcyto and GFP content mixing index. The fusion and mixing indexes are presented as means ± SEM of three independent experiments. Total number of nuclei counted in multinucleated cells and in cells in contact N ≥ 1000 for each experimental condition. Used unpaired t-test comparing each color (RFPcyto, GFP, mixed) for EFF-1 and HAP2 to the negative control (RFPcyto+GFP). * p<0.01; ** p<0.005; *** p<0.001; **** p<0.0005. **(D)** Still images from time-lapse experiments reveal merging of two mononucleated (i) and three (ii) cells expressing RFPcyto and HAP2 (arrows and arrowheads). Time indicated in h:min (see **Video** 1 and 2 for i and ii respectively). Note that the top nucleus (arrow in i) disappears due to defocus at 2:34. Two nuclei are out of focus at 4:57 (ii; see **Fig. S1 A**). Scale bars, 20 μm.

We asked whether HAP2/GCS1 family members display any similarity to known fusogenic proteins. HAP2/GCS1 are type I membrane glycoproteins composed of an N-terminal signal peptide, a large ectodomain, and a C-terminal cytoplasmic tail (Fig. 2 A). The ectodomain contains a conserved 50 amino acid region (Pfam PF10699, H/G domain, Fig. 2 A) that is unique to the HAP2/GCS1 protein family (Wong and Johnson, 2010). In search of structural similarities, we filtered and compiled ectodomains from all members of the HAP2/GCS1 family and subjected this dataset to two homology detection and structure prediction algorithms, HHblits (Remmert et al., 2012; Soding, 2005), and LOMETS (Wu and Zhang, 2007). We detected structural similarity to ectodomains of the EFF-1 protein from *C. elegans* (Pérez-Vargas et al., 2014) and class II viral fusogenic glycoproteins (Harrison, 2008; Igonet and Rey, 2012; Kielian et al., 2010; Podbilewicz, 2014; White et al., 2008) from three viral families, Flaviviridae, Togaviridae and Bunyaviridae (**Fig. S2 A, B**). Both methods detected similarity to class II fusogens throughout the entire HAP2/GCS1 family, consistent with their expected role in gamete fusion in evolutionary distant species (**Fig. S2 C and D**). To generate structural models of AtHAP2 we used I-TASSER (Yang et al., 2015). The overall fold of the AtHAP2 ectodomain model showed the canonical architecture of class II fusogens with three domains: a β-barrel domain I (DI), a mostly β-stranded elongated domain (DII), and the Immunoglobulin (Ig)-like C2-set topology module (DIII), separated by a linker from DI (Fig. 2 B). Global protein structure comparisons between these proteins, as measured by Z-score (Holm et al., 2008) and TM-score (Wu et al., 2007), indicate high similarity, with scores typical for proteins belonging to the same fold (Fig. 2 C; **Fig. S3 A**). Structural models with similar architecture were also obtained for HAP2/GCS1 proteins from several species (**Fig. S3 C**). An unrooted tree inferred from a structural dissimilarity matrix shows the structural relationship between families (Fig. 2 D). Although amino acid sequence conservation between AtHAP2, somatic and viral class II fusogens is low, most β-strands of AtHAP2 ectodomain are conserved and arranged in the same way as in *C. elegans* EFF-1 and viral fusogens (Fig. 2 E) (Pérez-Vargas et al., 2014). AtHAP2 also shares conserved cysteine residues with class II fusogens, and a prominent loop at the tip of DII (the cd loop, Fig. 2 E). In viral fusogens this loop is highly hydrophobic and necessary for anchoring at the host cell membrane (Podbilewicz, 2014), whereas in EFF-1 it is negatively charged (Pérez-Vargas et al., 2014). Unlike EFF-1, the AtHAP2 predicted cd loop is hydrophobic and flanked by charged residues (**Fig. S3 D**). Two additional loops neighbour the AtHAP2 cd loop (Fig. 2). One of these loops (between the i and j β-strands) forms part of the conserved H/G domain, is hydrophobic and point mutations at equivalent residues of *Chlamydomonas* HAP2/GCS1 block trafficking to the plasma membrane (Liu et al., 2015).

**Figure 2.**
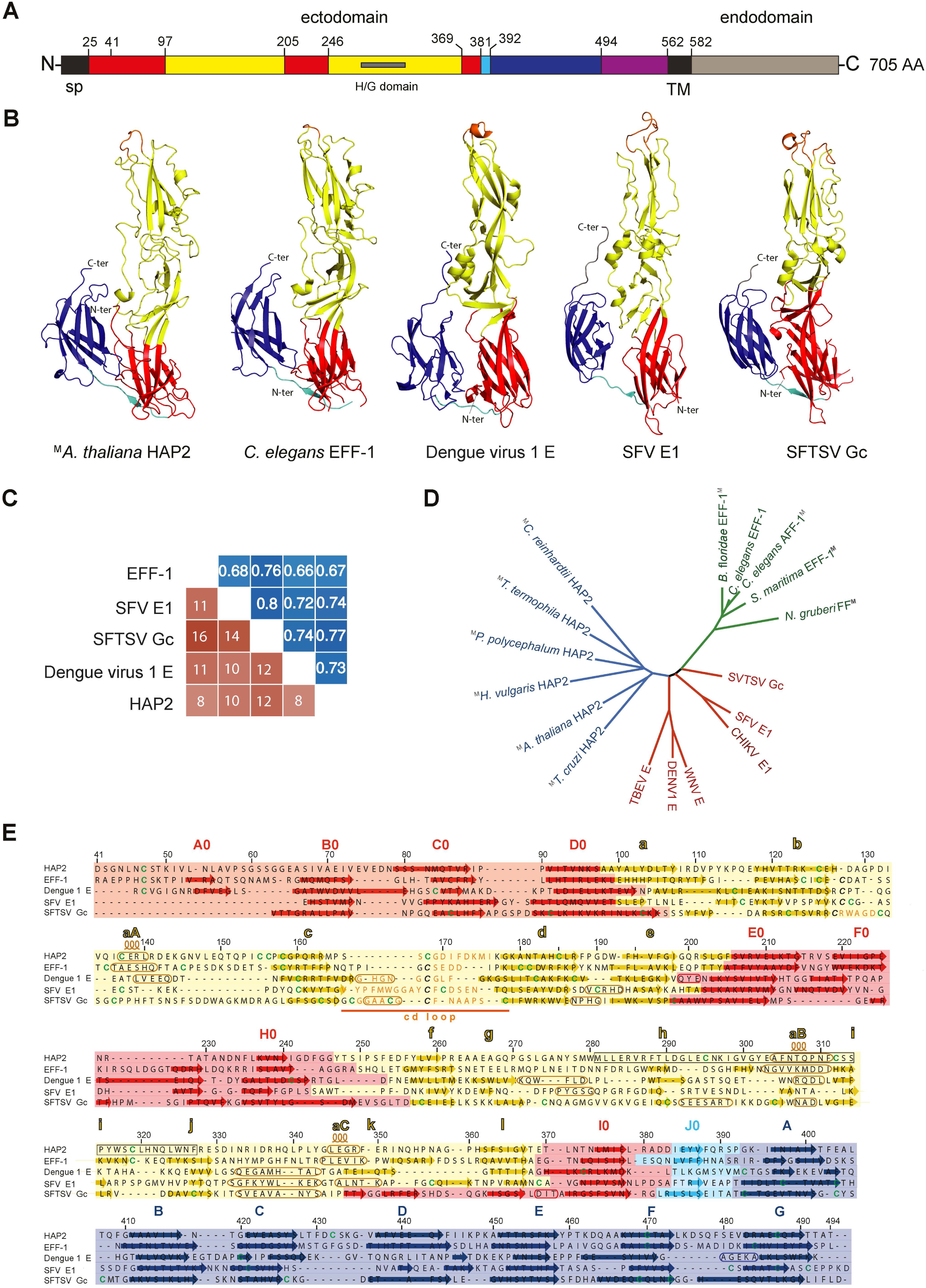
Model of the 3D structure of *Arabidopsis* HAP2 ectodomain indicates it is a class II fusogen. **(A)** Diagram of A. thaliana HAP2 protein coloured by domains according to the ectodomain modelled structure: signal peptide (sp) and transmembrane domain (TM), black; domain I, red; domain II, yellow, domain III, blue, domain I-III linker, cyan; stem, magenta and intracellular domain, grey. **(B)** Cartoon of AtHAP2 ectodomain modelled structure (residues 41-494), alongside with C. elegans EFF-1 (PDB 4ojc), Dengue virus E glycoprotein (DV1 E, PDB 4gsx), Semliki Forest Virus E1 glycoprotein (SFV E1, PDB 1rer) and Severe Fever with Thrombocytopenia Syndrome Virus glycoprotein Gc (SFTSV Gc, PDB 5g47) class II fusion proteins. Structures are coloured according to domains as in **A** and **E**; cd loops are shown in orange (facing up). **(C)** Structural similarity scores between experimental and modelled fusogens computed after flexible structural alignment. Blue matrix: TM-scores (values >0.5, are indicative of significant structural similarity), orange matrix: z-scores (values >2 are considered as indicative of significant structural similarity). **(D)** Unrooted tree inferred using a distance matrix. Boldface grey M indicates modelled structure, colors are HAP2, blue; EFF-1/AFF-1/FF, green; class II viral fusogens, red; see **Fig. S3 A. (E)** Structure-based alignment of AtHAP2 ectodomain with fusion proteins shown in **B**. Background colors indicate the domains organization (as in **A** and **B**), arrows in color and rounded rectangles denote beta sheets and alpha helices, respectively. The black box in AtHAP2 sequence marks the HAP2/GCS1 domain, the cd loops are marked in orange. Cysteines involved in conserved disulfide bonds stabilizing the cd loop are denoted in bold italics.

Class II fusion proteins mediate exoplasmic membrane fusion either unilaterally (e.g. from the virus envelope to the cell membrane) or bilaterally (e.g. EFF-1-mediated cell fusion (Podbilewicz, 2014; Podbilewicz et al., 2006)). HAP2 is essential in sperm for double fertilization in *Arabidopsis* (Johnson et al., 2004; Mori et al., 2006; von Besser et al., 2006). In *Chlamydomonas* and *Plasmodium* HAP2 is also active in male gametes (Liu et al., 2008). However, in the seven-sexed *Tetrahymena* efficient fertilization requires the presence of HAP2 in both fusing gametes (Cole et al., 2014). To test whether HAP2-mediated cell fusion requires HAP2 presence in either one or both fusing cells, we mixed HAP2-BHK-RFPcyto cells with BHK-GFP cells and determined the percentage of multinucleation in cells expressing RFPcyto, GFP or both. Whereas HAP2-BHK-RFPcyto cells had 35% ± 1.9 multinucleation, cytoplasmic content mixing between HAP2-BHK-RFPcyto and BHK-GFP cells was only 2% ± 0.3, undistinguishable from the negative control transfections (Fig. 3 A). These findings indicate that AtHAP2 is required in both cells for fusion to occur (Fig. 3 B).

**Figure 3.**
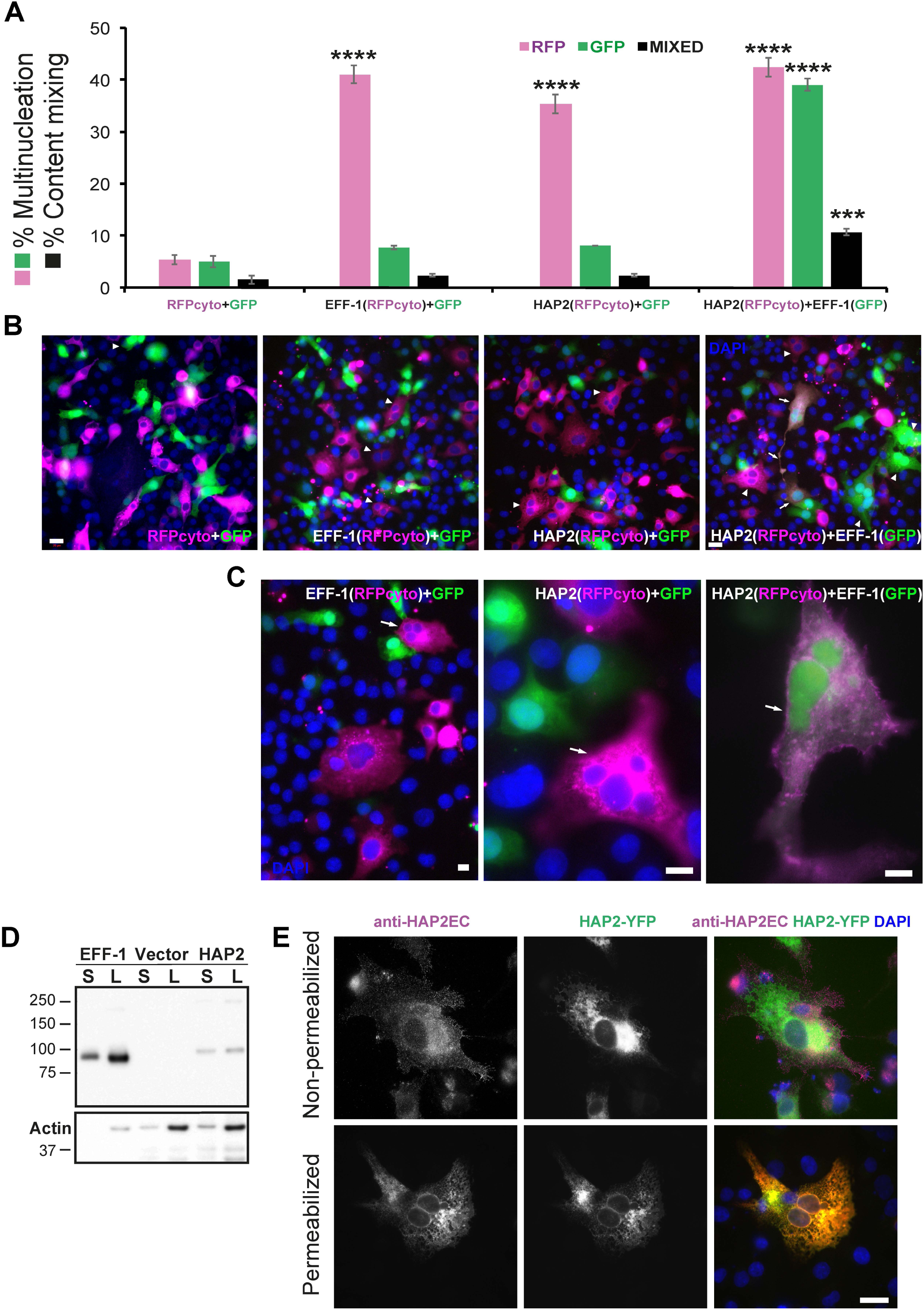
HAP2 is a bilateral fusogen. **(A)** Quantification of content mixing experiments showing that HAP2 and EFF-1 are required in both interacting cells to form hybrids. Control RFPcyto + GFP is the same as in Fig. 1 B. Total number of nuclei counted and unpaired Student’s t-test were performed as in Fig. 1 C. *** p<0.001; **** p<0.0005. Non-significant, p>0.05. **(B)** Representative fields used to determine percentages of multinucleation and content mixing. RFPcyto + GFP: Mixed control cells; Nuclear staining DAPI (blue). Dividing green cell with two nuclei (arrowhead). Scale bar, 20 μm. **EFF-1(RFPcyto) + GFP:** BHK-EFF-1 (magenta) do not mix with GFP-expressing cells, revealing that EFF-1-mediated fusion is bilateral (homotypic). Multinucleate cells expressing RFPcyto (arrowheads). **HAP2(RFPcyto) + GFP:** BHK-HAP2 multinucleation (arrowheads) and failure to mix with GFP-expressing cells revealing HAP2-mediated fusion is bilateral. **HAP2(RFPcyto) + EFF-1(GFP):** Hybrids (arrows) between BHK-EFF-1 (green) and BHK-HAP2 (magenta) reveal heterotypic merger of cells. Multinucleated green or red cells (arrowheads). Some hybrids are mononucleate probably due to cell division or nuclear fusion following merger. Scale bar, 20 μm. **(C)** Examples of multinucleate cells containing 2-4 nuclei are marked with arrows. The cells are able to divide after fusion therefore the number of nuclei per cells is usually smaller than 6. **EFF-1(RFPcyto) + GFP and HAP2(RFP) + GFP** images show no mixing indicating that HAP2-mediated fusion is bilateral (homotypic) in BHK cells. Scale bars, 10 μm. **(D)** Immunoblot of vector (negative control), EFF-1 and HAP2 proteins carrying a V5 epitope fused to the cytoplasmic tail. Surface biotinylation of BHK cells expressing the different proteins. S, surface expression after affinity purification using neutravidin agarose beads; L, lysate. The amount of sample of HAP2 “S” is 600 times higher than EFF-1. The amount of lysate for HAP2 “L” is 12 times higher than for EFF-1. **(E)** Immunofluorescence of HAP2-YFP shows surface expression in cells revealed by anti-HAP2Extracellular (HAP2EC) polyclonal antibody. Without permeabilization 20% of the cells expressing HAP2-YFP showed punctate expression on the surface; merged image (magenta). Untransfected BHK cells showed no immunoreactivity. Following permeabilization with detergent there is colocalization between immunostaining with anti-HAP2EC and the YFP signal revealing the specificity of the antibody that recognizes HAP2 in the reticular ER and perinuclear localization; merged image (yellow). Scale bar, 20 μm. Immunofluorescence in permeabilized BHK-HAP2-V5 cells using anti-V5 antibody shows similar localization (see **Fig. S1 B and C**).

We previously demonstrated that EFF-1 and its paralog AFF-1 can fuse BHK cells in a heterotypic way (Avinoam et al., 2011). To test whether AtHAP2 can promote cell fusion in trans with the nematode protein EFF-1, we mixed HAP2-BHK-RFPcyto cells with EFF-1-BHK-GFP cells and looked for cells containing both RFPcyto and GFP. We found that hybrids formed as efficiently as with EFF-1 in all fusing cells (10.7% ± 0.7; Fig. 3 A-C). Thus, EFF-1 can substitute for HAP2 in one of the cell membranes.

The extent of cell fusion is proportional to the level of surface expression of fusion proteins (e.g. EFF-1 and AFF-1) (Avinoam et al., 2011; Gattegno et al., 2007; Podbilewicz et al., 2006; Sapir et al., 2007). To determine whether AtHAP2 is expressed on the plasma membrane we used surface biotinylation followed by immunoblotting and immunofluorescence using an antibody against the extracellular domain of HAP2 in intact and permeabilized BHK cells. Following surface biotinylation we detected low amount of HAP2-V5 on the surface compared to EFF-1-V5 (Fig. 3 D). Immunofluorescence showed that AtHAP2 localized to puncta on the surface of 20% of the expressing BHK-AtHAP2-YFP cells (Fig. 3 E; **Fig. S1 B and C**). Since lower surface expression of AtHAP2, compared to EFF-1, resulted in similar fusogenic activity of both proteins, we conclude that AtHAP2 is a more efficient fusion protein compared to EFF-1.

Previously we demonstrated that eukaryotic fusogens can be assayed efficiently by replacing the Vesicular Stomatitis Virus (VSV) fusion glycoprotein G (VSVG) with a foreign fusogen and testing for the infectivity of the resulting pseudovirus (Avinoam et al., 2011). To test whether AtHAP2 can functionally substitute for VSVG, the fusogen glycoprotein G gene was deleted (VSVΔG) and replaced by HAP2 (VSVΔG-HAP2; Fig. 4 A). We first infected BHK-HAP2 cells with VSVΔG-G pseudoviruses and collected VSVΔG-HAP2 pseudoviruses from the supernatants. As controls we generated VSVΔG-EFF-1 particles (Avinoam et al., 2011). To test whether VSVΔG-HAP2 can infect cells we inoculated BHK cells, naïve or expressing HAP2 or EFF-1 (Fig. 4 B). We observed a 7 to 70-fold increase in infection efficiency in the fusogen-expressing cells compared to naïve cells (Fig. 4 C). Finally, to determine whether EFF-1 and HAP2 can interact through a bilateral mechanism in this virus-cell fusion system, we infected BHK-EFF-1 with VSVΔG-HAP2 and obtained similar infection efficiency to homotypic infection of BHK-HAP2. Likewise, VSVΔG-EFF-1 pseudotyped viruses showed similar infection efficiencies when infecting BHK-HAP2 or BHK-EFF-1 cells (Fig. 4 C). These finding reinforce our conclusion from cell-cell fusion assays (Fig. 3) and further suggest that HAP2 and EFF-1 directly interact to mediate heterotypic membrane fusion.

**Figure 4.**
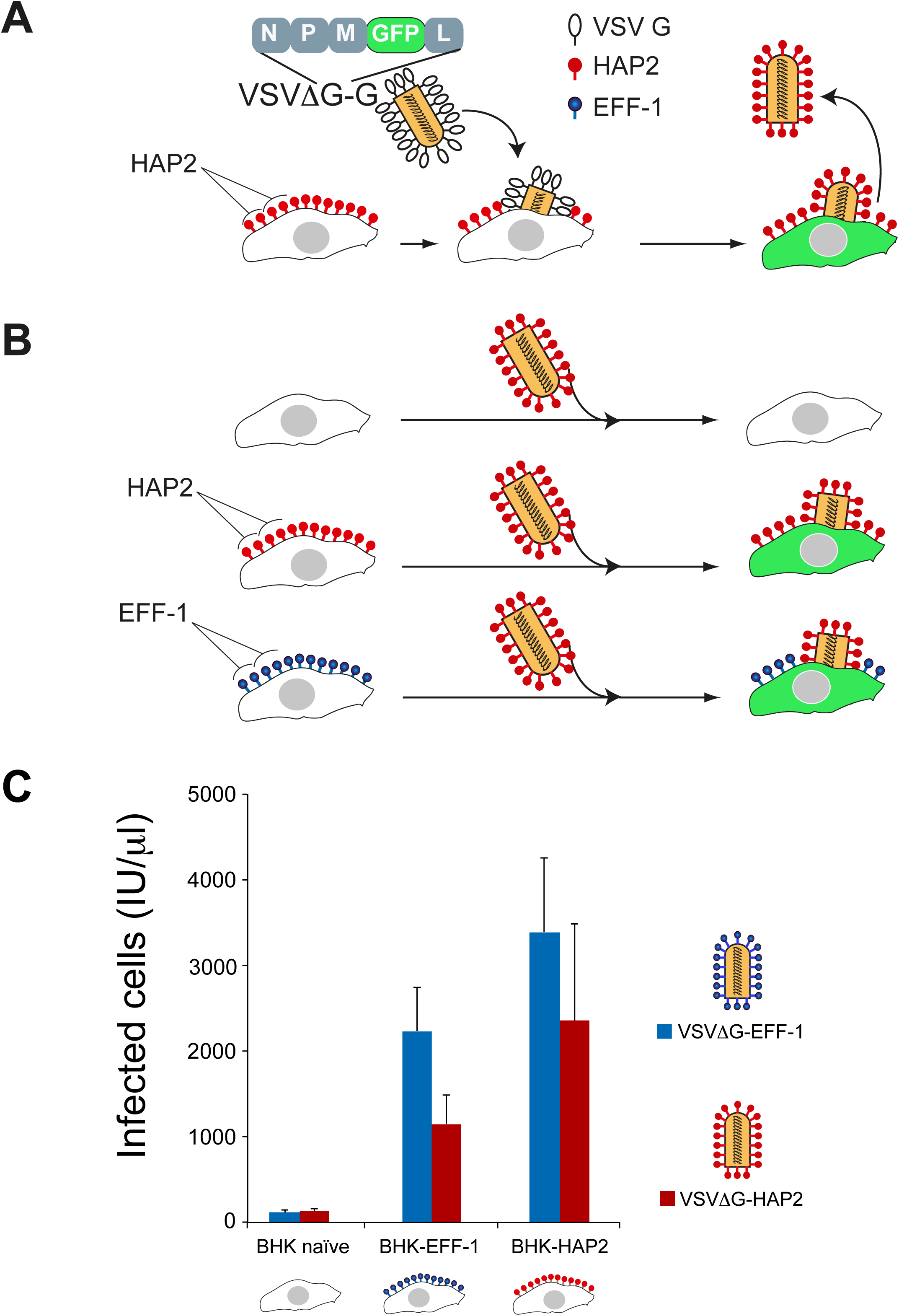
HAP2 can fuse the VSVΔG pseudovirus to target cells. **(A)** Cartoon illustrates the generation of VSVΔG-HAP2 pseudoviruses. Transfected BHK cells express HAP2 protein on the surface and were infected with G-complemented VSVΔG recombinant virus (VSVΔG-G). The viral genome encodes GFP replacing VSVG. Infection results in viral-induced expression of GFP by target cells (green cytoplasm). VSVΔG-HAP2 pseudoviruses were collected from the supernatant. **(B)** The activity of VSVΔG-HAP2 was tested on BHK (untransfected, naïve”), BHK-HAP2 and BHK-EFF-1 cells. Cartoon modified from (Avinoam et al., 2011). **(C)** Titers of VSVΔG pseudoviruses. The type of protein on the viral membrane (EFF-1 or HAP2) and on the BHK target cells (naïve, EFF-1 or HAP2) is indicated. Titers in infectious units (IU) represent the number of cells expressing GFP per microliter 24 h after virus inoculation. Data are mean ± SEM (n=3 experiments). We found no significant difference between infection with VSVΔG-HAP2 and VSVΔG-EFF-1 of the different BHK target cells (Two-tailed paired t test).

We then examined whether AtHAP2–mediated fusion proceeds via hemifusion, a fusion intermediate in which outer leaflets of two membranes are already merged but the inner leaflets remain distinct (Chernomordik and Kozlov,2005). To this end, cells transfected with AtHAP2 were labeled with both cell tracker as a cytosolic probe and DiI as a membrane probe, and were plated together with transfected but unlabeled cells. In this experimental design, cell fusion caught at a stage between hemifusion and fusion is expected to produce cells that acquired membrane- but not content-probe (Fig. 5 A). Indeed we observed the appearance of such cells (Fig. 5 B; arrows). Importantly, the number of cells labeled with only membrane probe was much lower in experiments in which the labeled cells expressing AtHAP2 were co-plated with unlabeled and non-transfected cells (Fig. 5 C). The apparent hemifusion in this control (~1.25%) could reflect background fusion or overlapping cells. Our results indicate that HAP2-mediated fusion proceeds through hemifusion intermediates. Our data are also the first evidence that a homotypic fusogen has to be present in both membranes to mediate even hemifusion, a fusion stage that is less energy demanding than opening and expansion of a fusion pore (Chernomordik and Kozlov, 2005).

**Figure 5.**
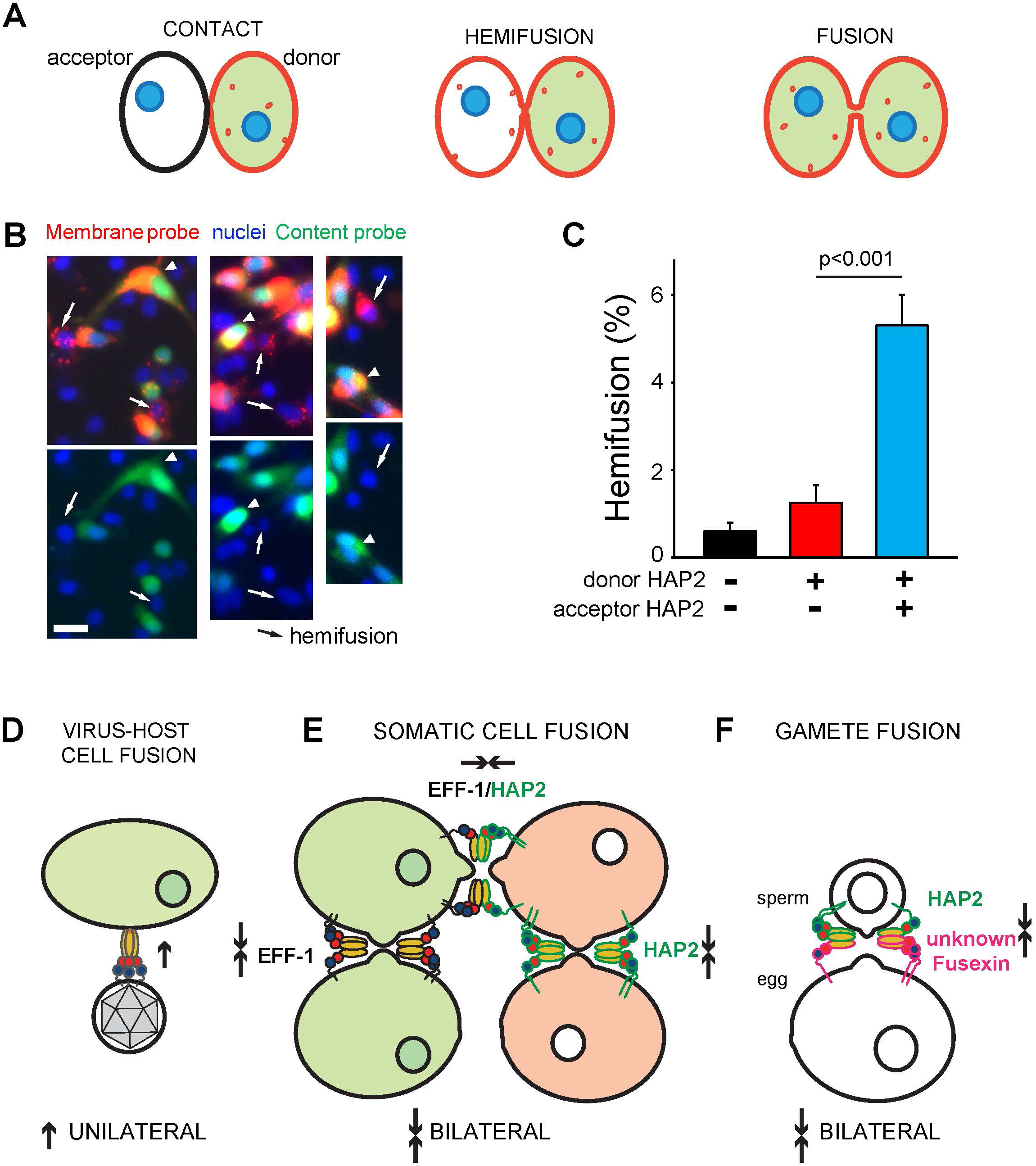
Hemifusion in HAP2-mediated fusion is bilateral and Fusexins use divergent mechanisms. **(A)** Cartoon illustrates the hemifusion assay where one cell type, labeled with two fluorescent probes, acts as the “donor” cell and the unlabeled cell acts as “acceptor”. Due to internalization of fluorescent lipid from the plasma membrane, by the time we score fusion this probe mostly labels intracellular membranes. **(B)** Fluorescence microscopy images of AtHAP2 transfected cells labeled with both cell tracker (green, content probe) and Vybrant DiI (red, membrane probe) co-plated with unlabeled AtHAP2 transfected cells. Top panel: green cell tracker, DiI (red), Hoechst 33342 (blue). Bottom panel: green cell tracker and Hoechst 33342 (blue). Hemifusion event is detected as an appearance of a cell (marked by an arrow) that acquired only membrane probe apparently from an adjacent double-labeled cell (arrowhead). Scale bar, 20 μm. **(C)** Hemifusion extents quantified as the ratio between numbers of cells labeled only with membrane probe and numbers of cells labeled with both membrane and content probes. The results are means± SEM (*n*=3). **(D)** Viral class II trimeric fusion proteins (viral Fusexins) have a unilateral fusion mechanism and the Fusexin is present only in the virus’ envelope or in one cell during cell-cell fusion. **(E)** Somatic cellular Fusexins (e.g. EFF-1 and AFF-1; on green cells) use a bilateral homotypic mechanism and the model proposes Fusexin has to be in both cells forming trans-trimeric complexes (Pérez-Vargas et al., 2014; Podbilewicz et al., 2006). Our results suggest a mechanistic model where sperm AtHAP2 fuses animal cells using a homotypic design (bilateral; red cells). HAP2 and EFF-1 can fuse cells in trans using a heterotypic mechanism. **(F)** We hypothesize that the egg of Arabidopsis (the central cell too) expresses an unidentified Fusexin that interacts with sperm HAP2.

Together, our findings indicate a somatic cell-like bilateral mechanism for AtHAP2 rather than a unilateral viral-like mechanism. Such bilateral mechanism is likely to operate not only in flowering plants but also in other organisms expressing members of the HAP2/GCS1 family (Fig. 5 D, E and F).

The observed structural and functional similarities between the gamete HAP2/GCS1 proteins, EFF-1 fusogen and class II viral fusion proteins could result from either common ancestry or convergence. Our data suggest a common ancestry based on the following observations: first, HAP2, EFF-1 and class II viral fusion proteins share a common domain architecture (Fig. 2). Second, proteins of the same fold are likely to be homologues with only a small fraction having different ancestries (Forslund et al., 2008). Third, comparisons of HMMs-based profiles of the complete ectodomain from different members of class II viral fusogens, HAP2 and FF (EFF-1/AFF-1) show low but significant amino acid sequence similarity between members of different groups (**Fig. S3 B**). Altogether, the combined conservation of structure, sequence and function makes a robust case for considering these proteins as diverged from a common ancestor.

Based on the ancient origin and function of exoplasmic membrane fusion we suggest naming this superfamily of fusion proteins the FUSEXINS (FUsion proteins essential for SEXual reproduction and EXoplasmic merger of plasma membranes). In conclusion, we have demonstrated the fusogenic activity of a fertilization protein and found a molecular link unifying sexual, somatic and viral membrane fusion. We hypothesize that Fusexins are ancient proteins and their emergence –either in viruses or cells— may have been a key innovation spurring sexual reproduction, exoplasmic somatic membrane fusion, and the rise of eukaryotes more than 2.5 billion years ago.

## Materials and Methods

### Homology detection

To detect structural homology in the HAP2/GCS1 sequences annotated in the InterPro database were taken as the HAP2 dataset (IPR018928). A total of 299 amino acid sequences were analyzed with TOPCONS (Tsirigos et al., 2015) to predict membrane topology. The resulting 299 ectodomain amino acid sequences were then used as queries against HHblits (Remmert et al., 2012) and LOMETS (Wu and Zhang, 2007). The Uniprot20 database (available on the Söding group server) was used to build Hidden Markov Models (HMMs) with one iteration of HHblits using default parameters. A subsequent search was performed using these HMMs against the PDB70 database (also available on the Söding group server) with default parameters to generate the final structural prediction using HMM-HMM alignment. To collect the results of the HHblits pipeline on all sequences, the output files were parsed with a script using CSB toolbox (Kalev et al., 2012). LOMETS, a meta algorithm that is part of the I-TASSER suite (Yang et al., 2015), was also used to predict structural homology for the 299 HAP2/GCS1 ectodomains. The standalone version uses six structural threading algorithms which are all variations of MUSTER (Wu and Zhang, 2008). LOMETS was run with default parameters for each HAP2 ectodomain sequence. LOMETS output files were parsed with a custom parser written in python to compile alignment template codes, columns aligned and z-scores (online supplemental material). All structural predictions using LOMETS and HHblits were performed locally. TOPCONS topology predictions were run on the TOPCONS server. Complete lists of structural prediction results are available in **Table S2**.

### Models Generation

Structural models were generated for a taxonomically diverse subset of ectodomain sequences using the I-TASSER server (Yang et al., 2015). EFF-1 and HAP2 ectodomains were prepared by removing signal sequences, transmembrane domains and C-terminal intracellular domains in the same way as for the structural prediction pipelines. Models were built from the ectodomains of sequences from the HAP2/GCS1 family available in the Uniprot database for *Arabidopsis thaliana* (F4JP36), *Hydra vulgaris (A3FEQ2), Physarium polycephalum* (Q2PGG5), *Chlamydomonas reinhardtii (A4GRC6), Trypanosoma brucei gambiense* (D0A4L4) and *Tetrahymena thermophile* (A0A060A682). Additional models for the AFF-1/EFF-1 family were built from *Naegleria gruberi* (D2W008), *Strigamia maritima* (T1ISJ2), *Caenorhabditis elegans* (G5EGL9), and *Branchiostoma floridae* (C3YJ9) ectodomains. Structural constraints based on probable disulfide bonds were used as additional inputs on the I-TASSER server for subsequent models for the *Arabidopsis* HAP2 ectodomain. These bonds were inferred from residue conservation in MSAs of HAP2/GCS1 ectodomains and visual inspection of the preliminary model. I-TASSER was run a second time on the *Arabidopsis* ectodomain sequence using these structural constraints to generate the final model presented in this publication. All of the PDB models generated using I-TASSER are available in the online supplemental material.

### Structure-based multiple sequence alignment

Structural alignments were used to generate high quality MSAs and pairwise superpositions of structures for use with structural similarity metrics. FATCAT (Ye and Godzik, 2004) pairwise alignments were run on all pairs of monomers in the I-TASSER models and crystal structures. Using a custom parser of the FATCAT output, pairwise alignments were generated in Fasta format. These alignments were iteratively merged using Clustal omega (Sievers et al., 2011) so that in each merger at least one sequence was shared between the Fasta files being merged until one a global alignment containing at least one of each of the sequences for the structures being aligned is produced. These final Fasta format alignments contain multiple sequence alignment possibilities for the set structures fed to the algorithm. Since all of the possibilities of sequence alignment are relatively similar, the final version of the alignment is chosen in Jalview (Waterhouse et al., 2009) by manual inspection.

### Structural similarity and structural phylogeny

Structural similarity matrices were built using the Dali Z-score (Holm and Rosenstrom, 2010) and the TM-score (Zhang and Skolnick, 2005) using pairwise superpositions generated by FATCAT. It has been shown that Z-scores above 2 (Holm and Rosenstrom, 2010) and TM-scores over 0.5 (Xu and Zhang, 2010) are indicative of proteins belonging to the same Folds, usually homologous. TM-score matrix was transformed into a distance matrix with the metric dist = 1 – TM-score. This structure distance matrix was used to infer a phylogeny of ectodomains using FASTME (Lefort et al., 2015) on the default settings. The automation script made for the generation of structure based phylogenies is available in online supplemental material.

### Profile based phylogeny

A taxonomically diverse set of ectodomain sequences of fusogens were used as search queries for HHblits against the Uniprot20 database to generate profiles, as hidden Markov models (HHM Format) incorporating also secondary structure information. For model construction realignment greediness (mact) was set to 10^-1^ and HHblits was run for 3 iterations on otherwise default parameters. These models were used in an all versus all search using HHsearch. Realignment greediness (mact) and secondary structure weight (ssw) in realignment in the All versus all HHsearch alignments were set to 10^-4^ and 3x10^-1^, respectively. The automation script made for the generation of profile based phylogenies is available in online supplemental material.

### Installation and configuration of structural similarity analysis and profile-to-profile tools

Python scripts used for structural and profile-to-profile analyses, are available in online supplemental material. They depend on SciPy, NumPy, Pandas, Biopython and the CSB toolbox python libraries. The structural phylogeny script requires the installation of TMalign (available on the Zhang lab server) and the PDB protein comparison tool (available on the PDB server). The profile-to-profile analysis script requires the installation of HHblits and HHsearch along with the uniprot20 and PDB70 databases (available on the Söding group server). These scripts were built for use in an Ubuntu 14.04LTS environment and have not been tested using other operating systems.

### Electrostatic calculations

To calculate electrostatic properties of the protein surfaces, the Adaptive Poisson-Boltzmann Solver (Baker et al., 2001) was used with default parameters, using atomic coordinates previously prepared with PDB2PQR (Dolinsky et al., 2007). Solvent-accessible surfaces were coloured by electrostatic potential and finally rendered in PyMol (https://sourceforge.net/projects/pymol/).

### Cells and reagents

Baby Hamster Kidney cells (BHK) were BHK-21 (ATCC). BHK cells were grown and maintained according to standard protocols using Dulbecco’s modified Eagle’s medium (DMEM), supplemented with 10% fetal bovine serum (FBS, Biological Industries, Kibbutz Beit Haemek, Israel), 100 U/ml penicillin and 100 μg/ml streptomycin (Biological Industries), 2 mM L-glutamine (Biological Industries), 1 mM sodium pyruvate (Gibco), 30 mM HEPES buffer pH 7.3 at 37⁰C (Biological Industries). Cells were grown at 37⁰ C with 5% CO_2_. Cells were transfected using a mixture of 8 μl Fugene HD (Promega) and 112 μl Opti-MEM (Gibco) with plasmids specified below in 1 ml for every 35 mm plate (Nunclone surface, Nunc) without replacing the medium (Avinoam et al., 2011).

### DNA constructs

The *Arabidopsis* HAP2 plasmids were derived from AtHAP2promoter::HAP2cds::YFP (PGL290; GenBank AF234315) (von Besser et al., 2006) that fully rescues *hap2(-) Arabidopsis* mutants (Wong et al., 2010). HAP2 ORF was amplified from PGL290 by PCR using primers 5’GCACTAGTATGGTGAACGCGATTTTAAT and TCCCGCGGACTCTCACGTAGTCTTTGTT. The amplified AtHAP2 fragment was subcloned into pIZT (Invitrogen) following SpeI and SacII digestion to obtain pIZT::AtHAP2-V5-HIS used for expression in insect cells. AtHAP2-V5 was amplified from pIZT::AtHAP2-V5-HIS using 5’ CGTTATCGATATGGTGAACGCGATTTTAAT and GCCCGGGTCACGTAGAATCGAGACCGA. The AtHAP2-V5 fragment was subcloned into pCAGGS vector using ClaI and XmaI to obtain pCAGGS::AtHAP2-V5. To subclone AtHAP2-V5 into pGENE B inducible system, pIZT::AtHAP2-V5 was PCR amplified using 5’GAGGTACCATGGTGAACGCGATTTTAAT and CCACTAGTTCACGTAGAATCGAGACCGA primers and cloned into pGENE B between the KpnI and SpeI restriction sites to obtain pGENE B::AtHAP2-V5. AtHAP2-YFP was subcloned into pCAGGS between the restriction sites ClaI and XmaI. pGL290 was used as template and the insert was PCR amplified using primers 5’CGTTATCGATATGGTGAACGCGATTTTAAT and GCCCGGGTTACTTGTACAGCTCGTCCA. pGENE B::AtHAP2::YFP was obtained by PCR amplification of PGL290 using primers 5’GAGGTACCATGGTGAACGCGATTTTAAT and CCACTAGTTCACGTAGAATCGAGACCGA. The fragment was cloned into pGENE B following digestion with KpnI and SpeI.EFF-1A::V5 was PCR amplified from pIZT:EFF-1A::V5 (Podbilewicz et al., 2006) using primers OR54 (TTAATTGGTACCACTATGGAACCGCCGTTTGAGTGG) and OR55 (AATTAAGCTAGCTCAACCGGTACGCGTAGAATCGAGACC) and cloned into pGENE B using KpnI and SpeI.

For mifepristone-inducible expression in BHK cells we used the GeneSwitch System (Wang et al., 1994) as recommended by Invitrogen (Avinoam et al., 2011). Cells were transfected with pGENE B::EFF-1A-V5, pGENE B::AtHAP2-V5 or pGENE B::AtHAP2::YFP. For transient expression of proteins we used pCAGGS::EFF-1-V5 (Avinoam et al., 2011), pCAGGS::AtHAP2-V5 or pCAGGS::AtHAP2-YFP.

To label nuclei and cytoplasm in mixing assays we used peGFP-N1 (Clontech) (Addgene plasmid cat #6085-1) kindly provided by Maya Haus-Cohen and Yoram Reiter. To label the cytoplasm we used pRFPnes (Hu et al., 2003) (pRFPcyto), kindly provided by Claudio Giraudo and Jim Rothman. Oligonucleotides were synthesized by IDT Sintezza, restriction enzymes were purchased from New England Biolabs (NEB) or Fermentas, and ligations were performed using T4 ligase (Promega).

### Fusion assays

For content mixing (fusion) assay, BHK cells at 70% confluence were co-transfected (using Fugene HD; Promega) with 2μg of pGENE B::EFF-1-V5 (Pérez-Vargas et al., 2014), or pGENE B::AtHAP2-V5; 1 μg of pSWITCH to obtain BHK-EFF-1 or BHK-HAP2 cells. As cotransfection markers we used RFP containing a nuclear export signal (pRFPcyto (Avinoam et al., 2011)) or peGFP-N1 (Clontech). Control cells were transfected with pRFPcyto or peGFP-N1. Four hours after transfection the cells were washed twice with PBS^-^ in the plates (35mm Nunc Surface) and detached using 0.05% EDTA solution (Biological Industries). BHK cells were collected in Eppendorf tubes, and resuspended in DMEM with 10% FBS, counted; equal amounts of cells were mixed and seeded in glass bottom plates (12 Well Black, Glass Bottom #1.5H; In Vitro Scientific). Four hours after mixing the EFF- 1 or AtHAP2 expression vectors were induced changing the medium with DMEM containing 10% FBS and 10^-7^ M mifepristone. After 18 h the cells were fixed with 4% paraformaldehyde in PBS and processed for immunofluorescence (see below). To assay multinucleated cells and to detect the transfected proteins (AtHAP2-V5 or EFF-1-V5), we stained cells with anti-V5 monoclonal antibody (Invitrogen) and nuclei with 1 μg/ml DAPI (Podbilewicz et al., 2006). Micrographs were obtained using Wide Field laser illumination using a Zeiss ELYRA system S.1 microscope at 20X (Plan-Apochromat 20X/NA: 0.8) or 25X magnification (LCI Plan-Neofluar 25X/ NA: 0.8). We counted the number of nuclei in cells expressing RFPcyto or GFP. For cytoplasmic mixing we counted the number of GFP nuclei in RFPcyto positive cells. Transfection efficiency was evaluated as 13-16% based on RFPcyto, GFP and anti-V5 immunofluorescence. The fusion indexes (shown as percentage of multinucleation) were defined as the ratio between the number of nuclei in multinucleated cells (Nm) and the total number of nuclei in multinucleated cells and expressing cells that were in contact (Nc) but did not fuse using the following equation:

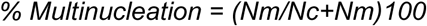

The percentage of multinucleation was calculated independently for GFP or RFPcyto expressing cells. DAPI was used to confirm the number of nuclei in expressing cells. GFP (also green nuclei) and RFPcyto (nuclei not red; “black”).

The GFP+RFP mixing index was calculated using the following equation:

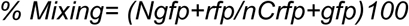

*Ngfp+rfp* is the number of GFP nuclei in cells with cytoplasmic RFP. *nCrfp+gfp* is the number of contacts between RFPcyto and GFP expressing cells. Note that RFPcyto or GFP mononucleated cells that were not in contact in the micrographs were not counted. The multinucleation and mixing indexes are presented as means ± SEM of three independent experiments (N ≥ 400 for RFPcyto, GFP and RFPcyto+GFP calculations in each experiment). The images were obtained using wide field microscopy with laser illumination in a Zeiss ELYRA system S.1 microscope at 63X magnification (Plan-Apochromat NA: 1.4).

### Hemifusion assay

BHK cells were grown in 35 mm tissue culture dishes to 75% confluence and transfected to express HAP2 as described above. 15 h later the cells were labeled with fluorescent lipid Vybrant DiI and CellTracker green (catalogue # V22885, and # C7025, ThermoFisher Sci.) at final concentrations of 4 μM and 1 μM, respectively (donor cells). 1 h later (at 16 h post transfection) the cells were co-plated with either unlabeled transfected cells or with unlabeled non-transfected cells at 1-to-1 ratio (acceptor cells). The cells were fixed with phosphate-buffered 4% formaldehyde for 10 min at 22°C and, 8 h later, labeled with Hoechst 33342 and analyzed using fluorescence microscopy. To quantify hemifusion, the numbers of cells labeled only with the membrane probe in each field of view were normalized to the total number of cells labeled with both membrane and content probes. For transfected donor cells mixed with transfected acceptor cells, the total number of double labeled cells was 1451. For transfected donor cells that were mixed with non-transfected acceptor cells the total number of double labeled cells analyzed was 1034. For non-transfected donor cells that were mixed with non-transfected acceptor cells the total number of double labeled cells analyzed was 1742.

### Immunofluorescence

BHK cells were grown on tissue culture 35 mm plates with glass coverslips or in glass bottom plates (12 Well Black, Glass Bottom #1.5H; In Vitro Scientific). Non-permeabilized cells were placed on ice and fixed with 4% paraformaldehyde in PBS, washed twice with ice cold PBS followed by incubation in 40 mM NH_4_Cl to block free aldehydes and blocked in ice cold 1% FBS in PBS. Permeabilized cells were fixed with 4% paraformaldehyde in PBS, followed by incubation in 40 mM NH_4_Cl to block free aldehydes, washed in PBS, permeabilized in 0.1% tritonX-100 in PBS and blocked in 1% FBS in PBS. After fixation the plates/coverslips were incubated 1 h with mouse anti-V5 antibody (Invitrogen) 1:500 or with rabbit anti At-HAP2 1:250 in PBS. The secondary antibodies were goat anti-mouse coupled to Alexa 643 or goat anti-rabbit coupled to Alexa 568 (Molecular Probes/Invitrogen) diluted 1:500 in PBS. Nuclei were visualized with 1 μg/ml DAPI (Avinoam et al., 2011). Permeabilized and non-permeabilized BHK cells were used as negative controls and gave background staining. The images were obtained using wide field microscopy with laser illumination in a Zeiss ELYRA system S.1

### Live imaging of fusing cells

Time-lapse microscopy to identify fusing cells was performed using a spinning disc confocal microscope (Yokogawa CSU-X) with a Nikon Eclipse Ti and a Plan Apo 20X (NA:0.75) objective. Six well plate with glass bottom (In vitro Scientific) were incubated in a Oko lab CO_2_ and temperature controlled chamber at 37°C and 5% CO_2_. Cells were co-transfected with pGENE B::AtHAP2-V5, pSWITCH and pRFPcyto to obtain BHK-HAP2 expressing RFPcyto. Six to 18 h post-induction images in DIC and red channels were obtained every 2-3 min in different positions of the plate using high gain and minimum laser exposure. Images were captured with a iXon 3 EMCCD camera (Andor). Fusing cells were identified based on mixing of cytoplasms containing RFPcyto. Identified syncytia were imaged at higher resolution using Apo 60X (NA:1.4) objective. Confocal z-series were obtained to confirm the formation of multinucleated giant cells. Image analyses were done in Metamorph and Image J.

### Cell cycle inhibition

To test whether multinucleation was as a result of failure of cytokinesis, we used 5-fluoro-2’-deoxyuridine (FdUrd) treatment to arrest BHK cells at G1/S phase (Dijkwel et al., 1986). BHK-21 cells were grown on tissue culture plates with glass coverslips (Marienfeld 1.5H). The cells at 70% confluence were co-transfected (using Fugene HD) with 2 μg/μl pCAGGS::AtHAP2 DNA and 1 μg/μl pEGFP-N1. For negative control BHK-21 were transfected with 1 μg/μl pEGFP-N1. Four hours after transfection the medium was changed with fresh DMEM with 10% FBS and 2 mM FdUrd (Sigma). 16-18 h after FdUdr addition the cells were fixed with 4% PFA, permeabilized, stained with DAPI and incubated with mouse anti-V5, followed by anti-mouse ALEXA 568. We calculated multinucleation (see mixing assay) and compared the multinucleation index with control transfected cells without FdUrd. The average results of two independent experiments are shown. N>800 nuclei for each experiment.

### Surface biotinylation of proteins expressed on BHK cells

BHK cells at 70% confluence were transfected with 2μg pCAGGS, pCAGGS::EFF-1-V5 or pCAGGS::AtHAP2-V5. 18-24 h post-transfection cells were washed twice with ice cold PBS++ and labeled with EZ-Link Sulfo NHS-Biotin (Pierce) for 30 min on ice. Surface biotinylation was followed by 4 washes with ice cold PBS++ and one wash with DMEM with 10% FBS to quench residual biotin followed by 2 washes with PBS++. After the washes, 300 μl of lysis buffer (50 mM Tris-HCl pH 8.0, 100 mM NaCl, 5 mM EDTA and 1% Triton X-100 supplemented with chymostatin, leupeptin, antipain and pepstatin and 10 mM Iodoacetamide) was added to each plate and the plates were incubated 15 minutes on ice. After removal of insoluble debris by centrifugation (10 min at 21,000 g), the lysate was mixed with NeutrAvidin Agarose Resin (Thermo) and SDS was supplemented to a final concentration of 0.3%. Affinity purification of biotinylated proteins was performed incubating for two to twelve hours at 4°C. The precipitated complex was then mixed with SDS-PAGE loading buffer with freshly added 50 mM DTT and incubated 5 minutes at 100°C. After pelleting by centrifugation the samples were separated on 4-12% SDS-PAGE gel (Bolt-Invitrogen) and analyzed by western blotting using anti-V5 mouse monoclonal antibodies (1:5000; Invitrogen) followed by 1:15000 HRP goat anti-mouse antibodies (Jackson). Loading was controlled using anti actin C4 monoclonal 1:2000 (MP, Biomedicals). Data shown are representative of at least three independent experiments (Podbilewicz et al., 2006).

### Pseudoviruses

We prepared pseudoviruses based on the method originally used to analyze Ebola virus glycoprotein (Takada et al., 1997) and modified for AFF-1 and EFF-1 (Avinoam et al., 2011). BHK cells at 70% confluence were transfected with 2μg pCAGGS::EFF-1-V5 or pCAGGS::AtHAP2-V5. After 24 h at 37°C in 5% CO2, cells were infected with VSVG-complemented VSVΔG recombinant virus (VSVΔG-G) at a multiplicity of infection (MOI) of 2-5 for 1 h at 37°C in 5% CO_2_ in serum free medium (DMEM). Following six washes to remove unabsorbed VSVΔG-G we incubated the cells for 24 h at 37°C. The supernatants containing VSVΔG-HAP2 and VSVΔG-EFF-1 were collected and centrifuged at 600g for 10 min at 4°C to clear cell debris. Virions were collected and concentrated by two consecutive centrifugations at 100,000 g through a 20%, and 10% sucrose cushion and resuspended in 130 mM NaCl, 25 mM Hepes pH 7.4.

### Determination of the titers of VSV pseudotype viruses

To determine the titer of each pseudovirus preparation, 10^4^ BHK cells were plated in each well of a 96 well tissue culture plate. BHK cells were transfected with 1 μg/ml DNA of pCAGGS::EFF-1-V5, pCAGGS::HAP2-V5 or non transfected. We performed six serial dilutions of the viruses that were used to inoculate the cells. Following 18-24 h of incubation, GFP expressing cells were counted in at least two dilutions using a Zeiss ELYRA system S.1 microscope at 20X (Plan-Apochromat 20X/NA: 0.8). Each experiment was repeated at least three times. All the infections were done in the presence of anti-G monoclonal antibody to prevent the activity of any residual VSVΔG-G (Avinoam et al., 2011).

### Data analysis

We did not predetermine sample size using statistical methods. The experiments were not randomized. We were not blinded to allocation during experiments and assessment of outcomes. Interobserver error was estimated for counting of multinucleated cells, cells in contact and content mixing experiments. To estimate inter-observer variation two investigators counted cells in contact, multinucleation and mixing of contents for the same 20 micrographs of different fields (examples of the micrographs used can be seen in Fig. 3 B). The differences between percentages of multinucleation and content mixing obtained by the two observers was less than 10%. Figures were prepared with Adobe Photoshop CS5, Adobe Illustrator CS6 and Image J.

### Statistical tests

The results are expressed as means ± SEM. For each experiment at least three independent biological repetitions were performed. We evaluated the significance of differences between mean values by using the unpaired t test function (Graph Pad software).

## Online supplemental material

Fig. S1 shows confocal sections of fused HAP2-BHK cells and low surface localization of AtHAP2 in mammalian cells. Fig. S2 shows graphic summaries of HHblits and LOMETS structural prediction results for proteins of the HAP2/GSC1 family. Fig. S3 shows global structural comparisons and HMMs profile to profile analyses of fusexins. Videos 1 to 4 show time lapse imaging of BHK-HAP2 cells undergoing cell fusion. Table S1 shows that FdUrd treatment increases HAP2-mediated cell fusion. Table S2 is a dataset including HHblits and LOMETS structural prediction results. ITASSERectodomains.pdb file contains the structures shown in Fig. S3 C. profilePhylogeny.py contains the automation script for generation of profile based phylogenies using HHblits and HHsearch. StructuresToMSAandPhylo.py contains the automation script for the generation of structure based phylogenies using TMalign and Fatcat.

## Acknowledgements

We thank Julian Wong and Mark Johnson for providing an anti-AtHAP2 ectodomain antibody and a clone containing the AtHAP2 coding sequence (pgl290). We thank Daniela Megrian, Agustina Olivera-Couto and Fernan Agüero for help with databases. Ofer Katz and Boaz Gildor for preliminary experiments and molecular biology. Liliana Avila Ospina for AtHAP2 subcloning and preliminary multinucleation assays. Eduard Baquero for advice on biochemistry. Rony Chanoch and Meytal Landau for preliminary structural bioinformatics studies. We thank Alejandro Colman-Lerner, Alberto Kornblihtt, Diego Alvarez, Ori Avinoam, Amir Sapir, Meital Oren-Suissa, Javier Matias Hernandez, Mark Johnson, Peter Walter, Gustavo Caetano-Anollés and Dan Cassel for helpful discussions and for critically reading the manuscript. This work was supported by the International Centre for Genetic Engineering and Biotechnology (grant CRP/URU11-01 to P.S.A.), the Fondo para la Investigación Científica y Tecnológica, Argentina (grant PICT-2014-3034, to P.S.A.), Israel Science Foundation, (grant 826/08, to B.P.), the European Research Council (Advanced grant ELEGANSFUSION 268843 to B.P.), and the United States-Israel Binational Science (grant 2013151 to L.V.C. and B.P.). The research in the L.V.C. laboratory was also supported by the Intramural Research Program of the Eunice Kennedy Shriver National Institute of Child Health and Human Development, National Institutes of Health. D.M. is a fellow from CONICET, Argentina. M.G. and H.R. acknowledge support from Agencia Nacional de Investigación e Innovación and Programa de Desarrollo de las Ciencias Básicas.

The authors declare no competing financial interests.

## Author contributions

B.P. and P.S.A. designed the experiments. C.V. designed and performed fusion assays and fluorescence microscopy experiments. D.M. performed most of the bioinformatics data gathering and data processing. E.L. and L.V.C. designed, performed and interpreted the hemifusion experiments. E.M. performed surface biotinylation. B.P. performed live imaging of cell fusion and counted some fusion assays. D.M., M.G., H.R. and P.S.A. designed the structural and evolutionary bionformatic strategies. C.V., D.M., H.R., M.G., B.P. and P.S.A. analyzed the data. P.S.A. and B.P. supervised the work and wrote the paper with input from all the other authors.

## Author Information

Correspondence and requests for materials should be addressed to P.S.A. and B.P.

## Abbreviations used

AtHAP2: *Arabidopsis thaliana* HAP2
BHK: Baby Hamster Kidney cells
EFF-1: Epithelial Fusion Failure 2
FUSEXINS: FUSion proteins essential for sexual reproduction and EXoplasmic merger of plasma membranes
GCS1: Generative Cell Specific 1
HAP2: Happless 2
HMMs: hidden Markov Models
VSV: Vesicular Stomatitis Virus
VSVΔG: VSV pseudoviruses in which the glycoprotein G gene was deleted

